# Scalable embedding fusion with protein language models: insights from benchmarking text-integrated representations

**DOI:** 10.1101/2024.08.24.609531

**Authors:** Young Su Ko, Jonathan Parkinson, Wei Wang

## Abstract

Protein language models (pLMs) have become essential tools in computational biology, powering diverse applications from variant effect prediction to protein engineering. Central to their success is the use of pretrained embeddings–contextualized representations of amino acid sequences–which enable effective transfer learning, especially in data-scarce settings. However, recent studies have revealed that standard masked language modeling objectives often produce representations that are misaligned with the needs of downstream tasks. While scaling up model size improves performance in some cases, it does not universally yield better representations. In this study, we investigate two complementary strategies for improving pLM representations: (1) integrating text annotations through contrastive learning, and (2) combining multiple embeddings via embedding fusion. We benchmark six text-integrated pLMs (tpLMs) and three large-scale pLMs across six biologically diverse tasks, showing that no single model dominates across settings. Fusion of multiple tpLMs embeddings improves performance on most tasks but presents a computational bottleneck due to the combinatorial number of possible combinations. To overcome this, we introduce greedier forward selection, a linear-time algorithm that efficiently identifies near-optimal embedding subsets. We validate its utility through two case studies, homologous sequence recovery and protein-protein interaction prediction, demonstrating new state-of-the-art results in both. Our work highlights embedding fusion as a practical and scalable strategy for improving protein representations.

## Introduction

Protein language models (pLMs) such as ESM [1] have catalyzed progress across a wide range of protein modeling tasks, including variant effect prediction [2], structure prediction [1], and protein engineering [3,4]. At the core of these advances lies the ability of pLMs to generate rich, contextualized embeddings of amino acid sequences, vector representations that encode biological features learned during pretraining. These representations have become fundamental inputs for downstream applications, powering everything from sequence classification models to structure-aware graph networks. This strategy allows researchers to offload much of the representational burden onto the pretrained model, which is especially valuable in low-data regimes. This strategy has proven effective in many settings [5–17], but recent work has revealed that the masked language modeling (MLM) objective of pLM pre-training are not always well-aligned with the representational needs of certain downstream tasks.

While scaling up pLMs might be expected to produce better, more generalizable representations, this trend does not always hold. Li et al. [18] showed that increasing ESM’s size from 43M to 650M parameters improved performance on secondary structure prediction, but yielded mixed returns on tasks involving protein function and tasks where coevolutionary signals are less informative. These findings suggest that representations learned through standard MLM pre-training may be misaligned with the requirements of many biologically meaningful tasks. This raises a central question: **how can we improve protein representations to better support diverse downstream applications, especially those that do not directly benefit from scaling**.

Addressing the root cause by creating a new, well-aligned pre-training task is nontrivial. An alternative and promising direction is to leverage existing, but often ignored, protein annotations. Recent studies such as ProteinCLIP [19] and ProTrek [20] have introduced “text+protein” language models (tpLMs) [21], which integrate textual protein annotations (e.g., UniProt functional annotations [22]) into the model’s latent space. These models apply contrastive learning to align protein embeddings with corresponding text embeddings, producing representations that are often more informative than those from standalone pLMs. While these works demonstrated outperforming similar sized pLMs, such as the 650M parameter ESM2 or the 420M parameter ProtBert [23] in downstream tasks, they have not been compared to significantly larger pLMs such as ESM2 3B, Ankh, and ProtT5. Benchmarking against these scaled-up pLMs is essential for understanding whether tpLMs offer genuine representational gains, or whether their performance can be matched simply by increasing model size.

Another promising strategy for improving protein representations is to combine multiple pretrained embeddings, an ensemble learning strategy referred to as embedding fusion. Given the diversity of pLMs in architecture, scale, and training data [24], different models may capture complementary aspects of protein biology. While individual pLMs can perform well on specific tasks, no single model consistently excels across all biologically meaningful settings. In natural language processing, embedding fusion has been shown to improve performance in tasks such as hate speech detection [25–27]. In protein modeling, however, fusion has only been explored in isolated cases, such as protein function prediction [28] and post-translational modification prediction [29], and its broader utility remains unclear. We hypothesize that this underuse stems from the absence of systematic benchmarking, leaving open the question of whether embedding fusion can serve as a general-purpose strategy for improving pLM representations.

In this work, we address the central question of how protein language model (pLM) representations can be improved to better support downstream tasks. We present the results of an extensive benchmarking effort, in which we trained 1,980 deep learning models across six biologically diverse datasets. First, we benchmark six state-of-the-art (SoTA) tpLMs against three large SoTA pLMs across six diverse datasets to assess whether incorporating textual annotations yields more effective representations. Second, we explore embedding fusion as a strategy to combine the complementary strengths of pretrained models. While embedding fusion proves effective, we find that identifying the optimal combination of embeddings poses a major challenge: exhaustive search is computationally impractical.

To address this, we benchmark two classic feature selection algorithms alongside our proposed linear-time selection method for efficiently identifying high-performing embedding combinations. We find that our linear-time method offers significant speed-ups while maintaining competitive performance. Finally, we conduct case studies to demonstrate the practical utility of our approach. By applying embedding fusion with our selection algorithm to improve the representations used for two biologically meaningful tasks–protein-protein interaction prediction and homologous sequence recovery–we achieve new SoTA results in both tasks. Together, these findings introduce a practical and scalable strategy for improving protein representations, crucial for downstream tasks that are poorly aligned with current pLM pre-training.

## Results and Discussion

Our study is organized into four main components:

1. **tpLMs vs. large pLMs**: We evaluate six tpLMs against three large-scale pLMs across six diverse tasks **(Fig. 1A)**.
2. **Embedding fusion**: Using tpLMs, we benchmark embedding fusion and investigate its validity as a strategy to enhance representations **(Fig. 1B)**.
3. **Embedding selection**: We assess the effectiveness of two established algorithms and our proposed linear-time algorithm for embedding selection **(Fig. 1C)**
4. **Case studies**: We apply our findings to two representative challenges, homologous sequence recovery and protein-protein interaction prediction **(Fig. 1D)**.

**Figure 1.**
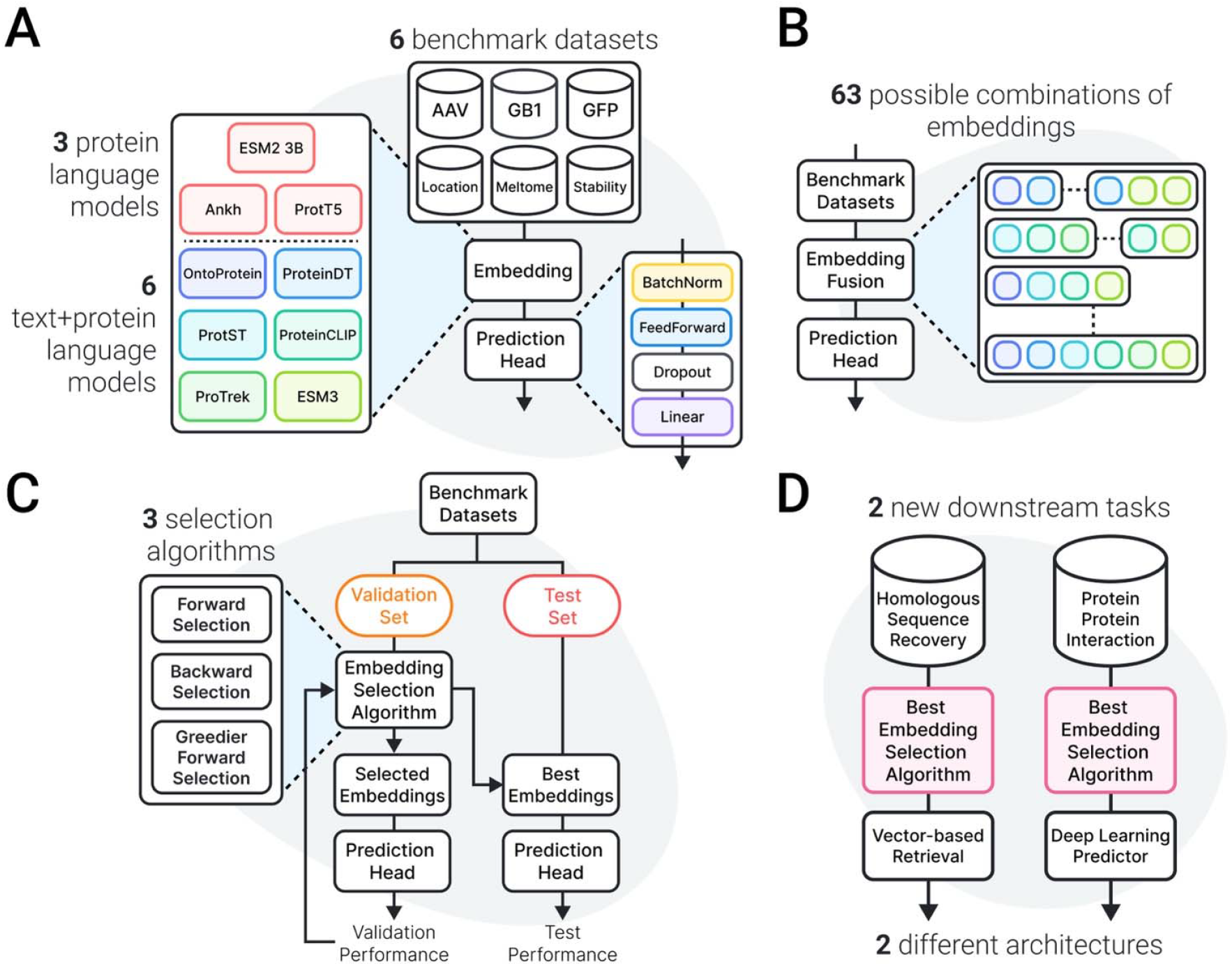
Study overview and benchmarking results. **A.** Six different text-integrated protein language models (OntoProtein, ProteinDT, ProtST, ProteinCLIP, ProTrek, and ESM3) are evaluated on six different datasets. Results are compared with ESM2 3B, Ankh, and ProtT5. **B**. For each dataset, all 63 possible combinations of text-integrated protein language model embeddings are benchmarked and compared with the best single tpLM embeddings. **C**. We benchmark forward selection, backward selection, and greedier forward selection in their ability to get as close to the global optimum on the same benchmark datasets **D**. We apply embedding fusion with greedier forward selection on two new tasks–homologous sequence recovery and protein-protein interaction prediction.

### tpLMs: promising, but not perfect

We first curated six SoTA tpLMs: OntoProtein, ProtST, ProteinDT, ProteinCLIP, ProTrek and ESM3 **(Fig. 2A)**. The selection criteria and key characteristics of each tpLM are further described in the **Methods and Supplementary Section S1**. To evaluate their effectiveness, we compared their representations against three large-scale pLMs, ESM2 3B, Ankh, and ProtT5, across six benchmark datasets (**see Methods and Supplementary Section S2**). Following Schmirler et al. [30], we used a standardized prediction head and adopted their training regimen, including model architecture and hyperparameters.

**Figure 2.**
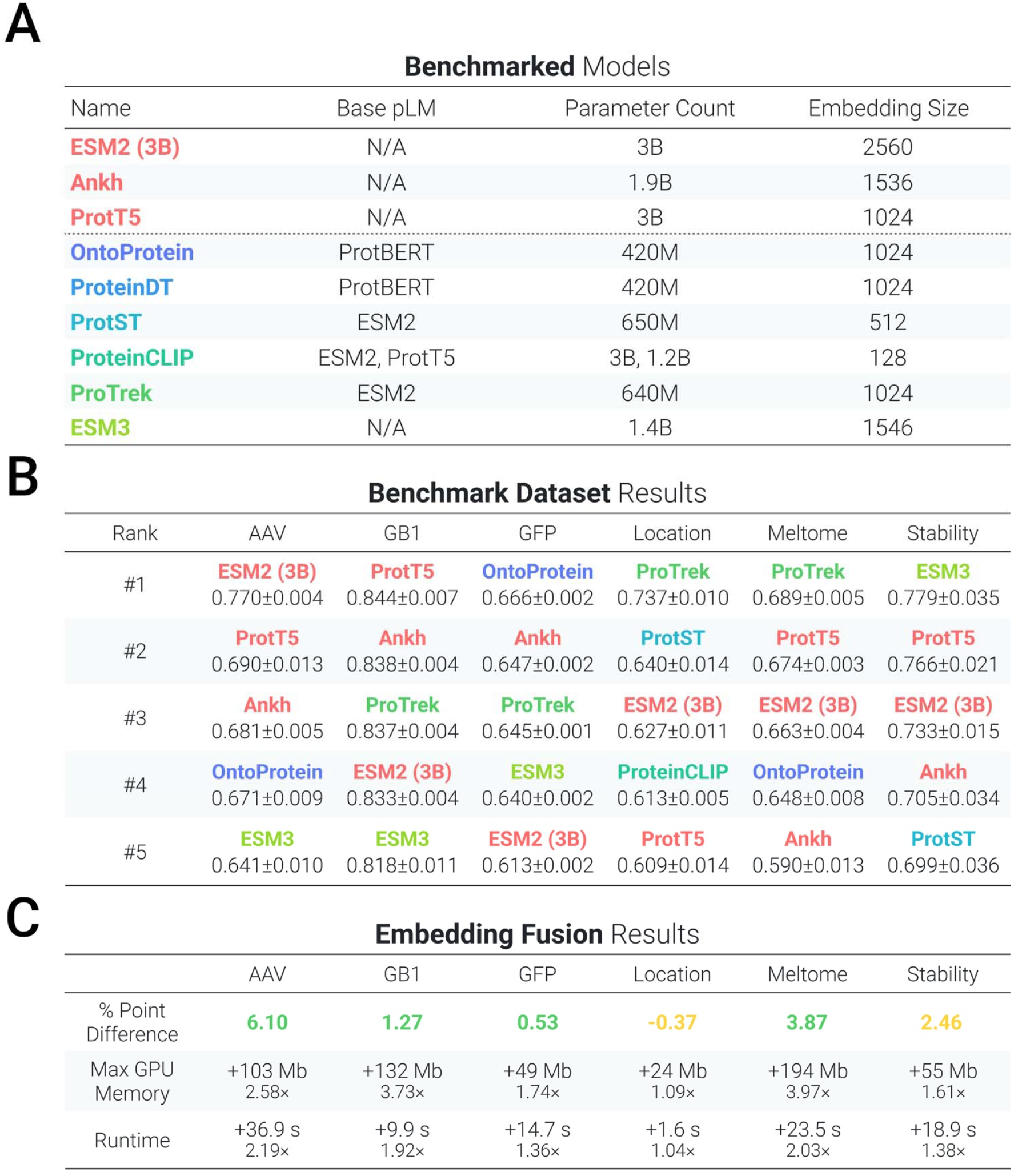
Results of benchmarking text-integrated protein language models and embedding fusion. **A.** Overview of protein language models benchmarked. **B**. Benchmarking results for text-integrated protein language models and text-free protein language models. 95% CIs are shown. **C**. Percent-point difference in performance gained by using embedding fusion compared to using a single text-integrated protein language model embedding. The difference in maximum GPU memory usage and runtime (absolute difference and fold-change) between the fused embedding and single embeddings are shown.

As shown in **Fig. 2B**, based on the average performance (accuracy for the Location benchmark, Spearman’s ρ for all others) on the respective test sets across five random seeds, a tpLM embedding achieves the best performance except for the AAV and GB1 benchmarks. Average performance of all models are shown in **Supplementary Table 1**. For GB1, ProTrek, the best tpLM, shares overlapping 95% confidence intervals (CIs) with ProtT5, the best model, indicating comparable performance. In the case of AAV, prior work has shown that performance continues to improve with larger model sizes, even when the task is poorly aligned with pre-training, suggesting this benchmark may benefit more from scale than representation alignment [18]. These results highlight both the promise and limitations of tpLMs: while text integration can improve representation quality, no single embedding consistently excels across all tasks. Notably, we observe the same trend for the large-scale pLMs–performance varies by task, with no pLM emerging as consistently superior. This variability led us to hypothesize that combining embeddings could combine their individual strengths to yield more robust and generalizable representations.

### Embedding fusion: effective but impractical

To investigate this hypothesis, we assessed every single possible combination of tpLM embeddings on the six benchmarks, using the same training setup as before (**Supplementary File 1**). For six tpLMs, 63 unique combinations are possible, where a combination is created by the concatenation of individual embeddings, disregarding order. While embeddings can be fused in different ways, at this stage, we establish a baseline with the most straight-forward approach. For each benchmark, we compared the embedding combination with the highest test performance with the single embedding with the highest test performance.

We observed that embedding fusion outperformed single embeddings in four out of six benchmarks, with statistically significant improvements in Spearman’s rank correlation coefficient (ρ) supported by non-overlapping 95% confidence intervals (**Fig. 2C**). Although no statistically significant improvement was observed for the Stability benchmark, a numerical increase in Spearman’s ρ was shown. For the Location benchmark, we observed a non-significant decrease in accuracy. Upon further investigation, ProTrek is the only tpLM to both update the text encoder and utilize the “Subcellular Location” annotations from UniProt during pre-training, suggesting its learned representations are already saturated for this task. Recognizing that embedding fusion increases the input dimension, and thus the parameter count in the projection layer, we additionally compare parameter-normalized performances **(Supplementary Table 2)** and observe the same trend of significance.

A possible concern is the increased resources necessary for embedding fusion. The best-performing combinations incurred an average 1.65× increase in runtime and 2.45× increase in peak GPU memory usage compared to the best single embedding. While these appear substantial as ratios, we emphasize that the absolute increases were modest, no more than 40 seconds in runtime or 200MB in memory. These multiplicative factors are inflated by the small size of our base model, where fixed overheads such as I/O and input projection dominate the runtime. As model size increases, the relative cost of fusion decreases, since the majority of computation shifts to deeper network layers. We demonstrate this empirically in **Supplementary Section S4**, where we show that fusion overhead does not scale linearly with model size. Thus, we reasoned these overheads are not prohibitively costly. Additionally, as Schmirler et al. [30] who ran evaluations on the same datasets with single embeddings reported, we observed differences in validation and test performance, which can be attributed to the individual dataset characteristics, such as dataset size or large domain shifts.

However, a crucial limitation is that the best combination of embeddings is dataset dependent and combining all embeddings does not lead to the best results, likely due to feature redundancy. This raises a practical challenge: how can one efficiently identify an optimal subset of embeddings without exhaustively evaluating all possibilities? Our current strategy evaluates all 63 combinations, which is computationally expensive and scales poorly as the number of available embeddings grows. To underscore the computational cost, note that ten embeddings yield 1,023 possible combinations, while fifteen create over 32,000. This combinatorial burden likely contributes to why embedding fusion remains underutilized in protein machine learning workflows, despite its clear performance benefits.

### Greedy algorithms enable tractable embedding fusion

The task of selecting an optimal set of embeddings can be framed as a feature selection problem, a well-studied challenge in classical machine learning in which the goal is to determine an optimal set of features. Greedy algorithms for feature selection such as forward selection and backward selection offer a tradeoff between performance and efficiency [31]: while they do not guarantee finding the global optimum, they can identify high-performing subsets at a fraction of the computational cost.

We adapt these strategies to the embedding fusion setting and evaluate three algorithms: forward selection, backward selection, and our proposed greedier forward selection (**Fig. 3A**). The design of greedier forward selection was motivated by two consistent trends in our benchmarking results: (1) the best-performing single embedding is nearly always included in the optimal combination, and (2) even the weakest individual embeddings occasionally contribute meaningfully to the fused representation. To balance these dynamics, greedier forward selection initializes from the top-ranked single embedding and incrementally adds others based on individual performance, evaluating combinations at each step. Full algorithmic details are provided in the **Methods**.

**Figure 3.**
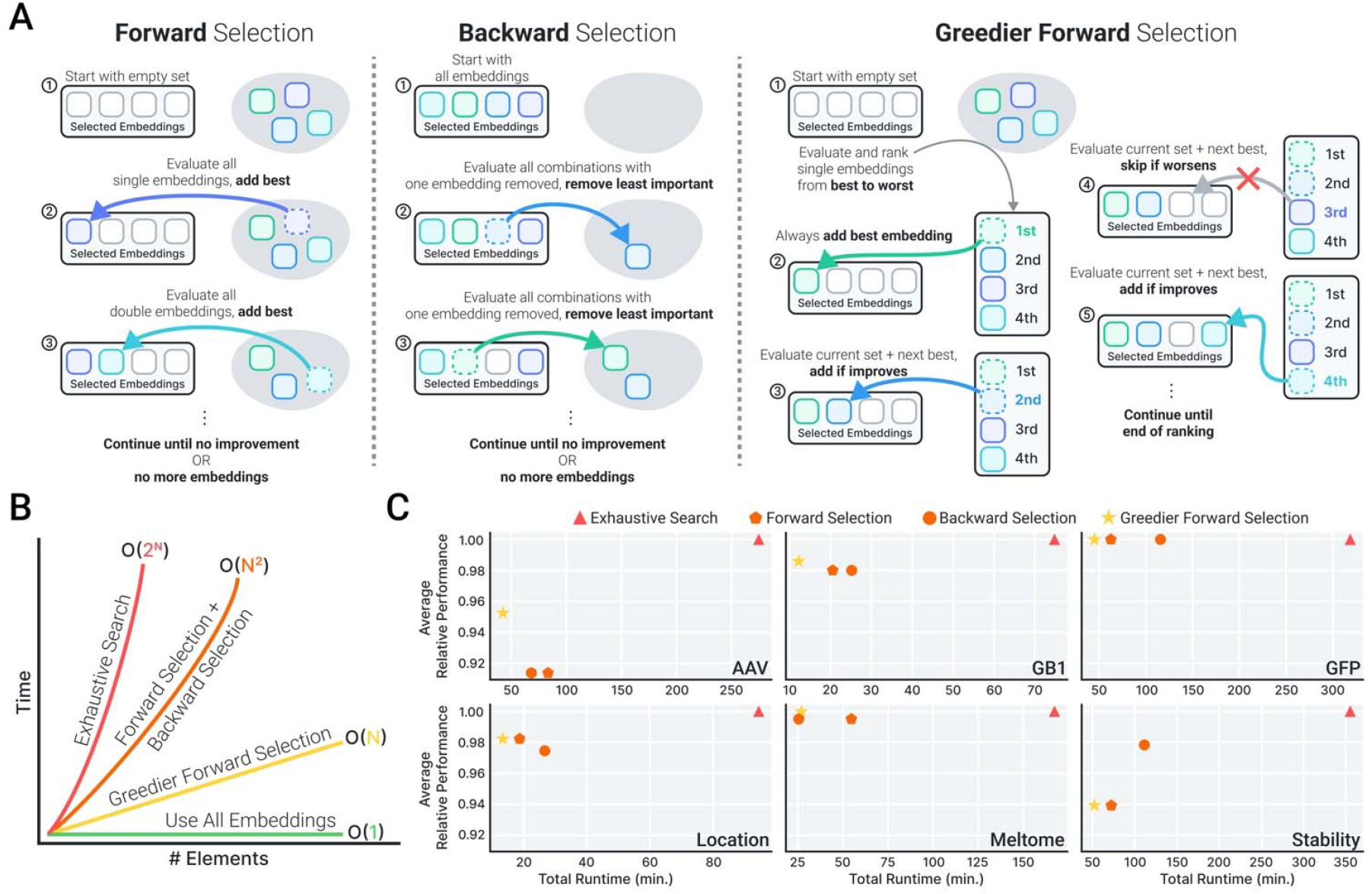
Benchmarking and analysis of embedding selection algorithms. **A.** Schematic overview of the embedding selection strategies evaluated: forward selection, backward selection, and greedier forward selection. **B**. Computational complexity of each selection method, visualized alongside baseline strategies–exhaustive search and naive use of all embeddings–to highlight scalability differences. **C**. Performance comparison across six benchmark tasks, showing how each selection method approximates the optimal result from exhaustive search.

As shown in **Fig. 3B**, all three greedy algorithms are theoretically more efficient than exhaustive search, which scales exponentially with the number of embeddings. In **Fig. 3C**, we benchmark each algorithm across all six datasets by comparing their performance to the best test performance obtained via exhaustive search. On average, greedier forward selection achieves 99.5% of the maximum test performance identified by exhaustive search, while being 1.69× faster than forward selection (which achieved 99.8%) and 2.11× faster than backward selection (99.2%). These results demonstrate that greedier forward selection strikes an effective balance between accuracy and efficiency, offering a scalable solution to embedding fusion that avoids combinatorial explosion.

We note that these benchmarks are designed to evaluate representation quality rather than to develop production-ready models. Our goal is to systematically test the utility of embedding fusion under controlled, reproducible conditions. Thus, as a final step in this study, we investigate how embedding fusion performs in such realistic, task-specific settings using existing state-of-the-art models.

### Case Study: Strengthening protein representations with embedding fusion

While our benchmarks were designed to systematically evaluate representation quality under controlled conditions, they do not fully capture how embedding fusion performs in real-world scenarios, where models are often fine-tuned, hyperparameter-optimized, or enhanced with techniques like LoRA. To assess whether the benefits of fusion persist in these practical settings, we conduct two case studies on biologically meaningful tasks: homologous sequence recovery and protein-protein interaction (PPI) prediction [19].

In each case, we apply embedding fusion–guided by greedier forward selection–to augment an existing state-of-the-art model. We fix all other components of the pipeline to isolate the contribution of improved representations. For the PPI task, we additionally compare against previously reported results from fine-tuning and LoRA-based approaches to contextualize our approach within the broader landscape of machine-learning strategies. Finally, we evaluate how close our heuristic selection gets to the global optimum by running exhaustive search. These case studies move beyond controlled benchmarking to demonstrate how fusion can enhance the performance of real, task-specific models, reinforcing its utility as a practical and accessible strategy.

### Embedding fusion enabled more accurate homologous sequence recovery

The homologous sequence recovery task evaluates protein embeddings in a nearest-neighbor retrieval setting, where predictions are made by comparing embeddings directly via cosine similarity. More specifically, a query protein embedding is compared against a database of embeddings using cosine similarity and retrieves the most similar embedding. A retrieval is considered correct if retrieved protein shares the same CATH [32] superfamily **(Fig. 4A)**. Unlike our deep learning benchmarks, this task requires no training or parameter tuning, offering a complementary perspective on representation quality. Additionally, the reduced computational burden enables evaluation across all pairwise combinations of the six tpLMs and three large pLMs.

**Figure 4.**
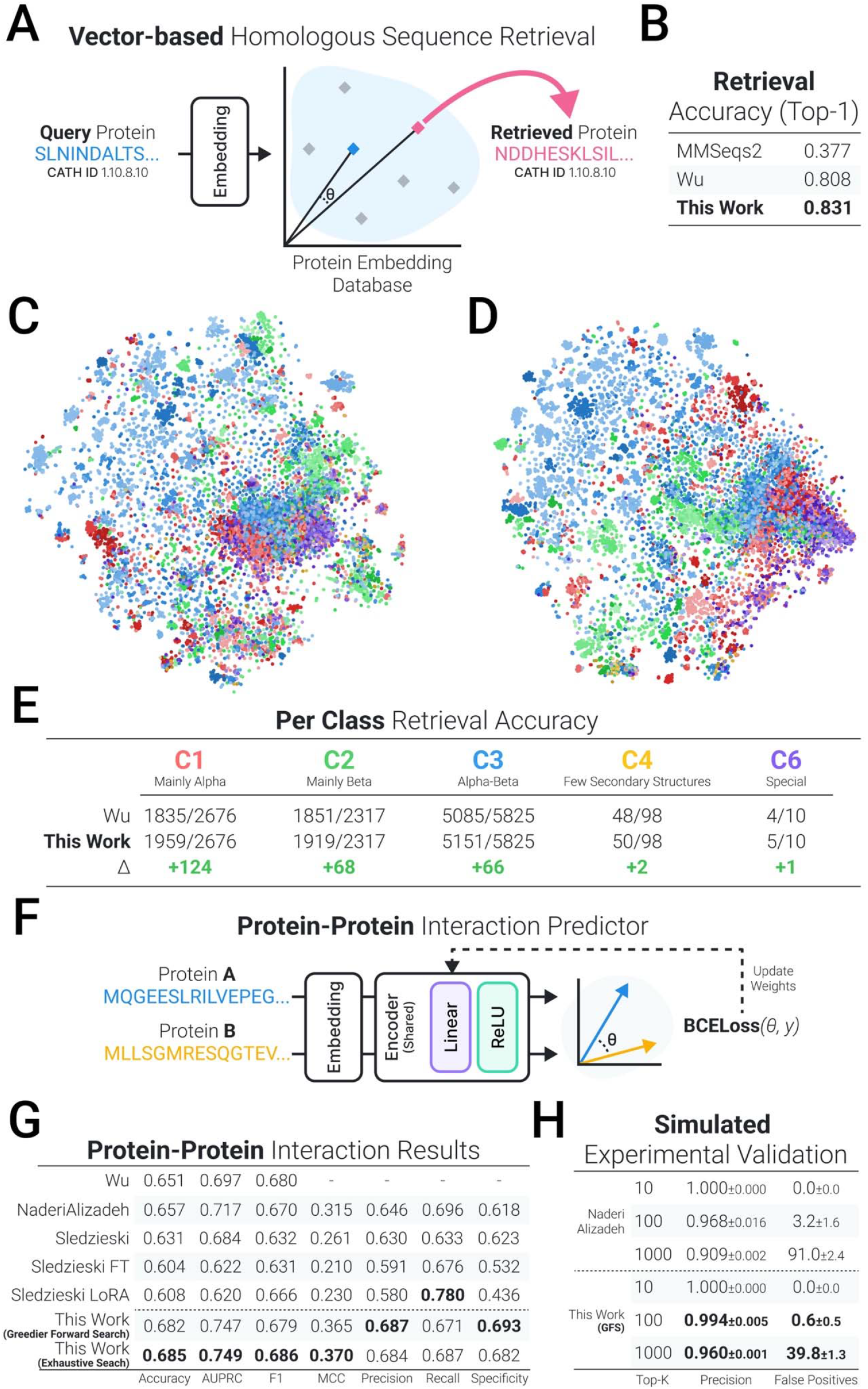
Case study design and outcomes for homologous sequence recovery and protein-protein interaction (PPI) prediction. **A.** Overview of the homologous sequence recovery task: embeddings are used to retrieve homologous domains based on cosine similarity. **B**. Top-1 accuracy across individual and fused embeddings for the recovery task, highlighting improvements from embedding fusion. **C**. t-SNE visualization of CATH domain embeddings using the previous state-of-the-art method (ProteinCLIP). **D**. t-SNE visualization of the fused embeddings selected via greedier forward selection, showing improved clustering of CATH classes. **E**. Comparison of per-class retrieval accuracy between ProteinCLIP and the selected fusion combination, illustrating class-specific performance gains. **F**. Performance of embedding fusion on the PPI prediction task. **G**. Precision and false positives for top-k ranked predictions, simulating experimental prioritization in a resource-limited setting.

The previous state-of-the-art was introduced by Wu et al., who used ProteinCLIP embeddings to achieve higher top-1 retrieval accuracy than MMseqs2 [33]. Using greedier forward selection, we identified a fused embedding composed of Ankh, ProtT5, ESM3, ProteinCLIP, and ProTrek that achieves an accuracy of 83.1%, surpassing the previous best of 80.8% **(Fig. 4B)**. Notably, this combination was also identified as the global optimum when compared against all possible combinations via exhaustive search.

To better understand how embedding fusion alters the structure of the representation space, we visualized all CATH sequences using t-SNE, comparing our fused embeddings **(Fig. 4C)** to the previously best-performing ProteinCLIP embeddings **(Fig. 4D)**. We used 3,000 iterations, cosine distance, and a perplexity of 50. The fused embeddings exhibit improved class separation and tighter intra-class clustering, suggesting that they better capture the distinct structural features of each CATH class. While t-SNE provides qualitative insight, we also sought a quantitative measure of improvement. We therefore computed per-class retrieval accuracies and observed consistent gains across all classes **(Fig. 4E)**, in agreement with the visual patterns.

These results demonstrate that embedding fusion, when guided by a lightweight selection strategy, can meaningfully enhance protein representations even in a training-free retrieval setting. The improvement over the previous state-of-the-art, combined with the ability of greedier forward selection to recover the global optimum, underscores the practical viability of fusion in real-world applications. Moreover, the qualitative and quantitative gains in class separation and retrieval accuracy suggest that fused embeddings capture richer and more discriminative structural features than any individual model alone. Notably, the best-performing combination includes both Ankh and ProtT5, illustrating that the benefits of embedding fusion extend beyond text-integrated models and generalize to text-free pLMs as well.

### A new frontier for sequence-based protein-protein interaction prediction

Our second case study returns to a deep learning setting, focusing on the widely studied challenge of sequence-based PPI prediction. Such models are essential for high-throughput in silico screening, rational binder design, and the discovery of novel molecular interactions. However, early approaches were often confounded by data leakage, where high sequence similarity between training and test sets inflated performance metrics [34]. To address this, Bernett et al. introduced a rigorously filtered benchmark of human PPIs that minimizes sequence redundancy and has since become the gold standard for evaluating generalizability in this task.

To examine how embedding fusion performs in this real-world, leakage-free setting, we compile results from recent literature, including models that leverage fine-tuning and LoRA on ESM2 [19,35,36]. We adopt the architecture and hyperparameters of NaderiAlizadeh and Singh [35], who report the highest accuracy on the Bernett benchmark (**Fig. 4F**; see **Methods** for full details). We modify only the embedding module, using greedier forward selection to identify the most effective combination of pretrained embeddings based on validation performance. This setup allows us to directly assess whether improved representations alone can enhance performance in an already-optimized model.

The selected combination, ProteinCLIP, ProtST, and ProTrek, achieved the highest average AUPRC and outperformed all individual tpLM embeddings **(Supplementary Table 3)**. On the test set, this combination achieves a new state-of-the-art in both AUPRC and accuracy (**Fig. 4G**). Exhaustive search revealed that the true best-performing combination includes an additional model, ProteinDT, further improving test AUPRC and pushing performance beyond previously reported baselines. Importantly, both the selected and global-optimum combinations outperform any single tpLM, demonstrating the consistent benefit of fusion in this task.

To better understand the practical implications of this improvement, we simulated a realistic use case in which a classifier is used to select top-ranked interaction candidates for experimental validation. We re-implemented the NaderiAlizadeh model, including its original use of ESM2 650M embeddings (see **Supplementary Section S5** for reproduction details) and compared it against the fusion-enhanced model. For each model, we identified the top-k predictions (k = 10, 100, 1000) from the test set with the highest predicted probability of interaction and calculated the precision and number of false positives at each threshold **(Fig. 4H)**.

Both models achieved perfect precision in the top-10 predictions. However, at broader thresholds, the fusion-based model demonstrated markedly better performance, achieving higher precision and fewer false positives and producing less than half as many false positives as the baseline. These findings highlight how improved representations from fusion can yield more robust classifiers, better able to generalize and prioritize meaningful interactions, especially in scenarios where false positives are costly.

## Concluding Remarks

This study set out to investigate how pLM representations can be improved, particularly for tasks that remain misaligned with current pre-training objectives. We first benchmarked six tpLMs against three large-scale pLMs across six diverse tasks. While tpLMs frequently outperformed larger models, no single model consistently yielded the best representations across all benchmarks. These results underscore a key limitation in current approaches: relying on a single model, regardless of size or supervision, does not capture the full spectrum of biologically relevant features.

To address this, we explored embedding fusion as a general-purpose strategy for combining complementary representations. While fusion consistently improved downstream performance, its adoption has been limited by the computational burden of exhaustively evaluating all embedding combinations. To make embedding fusion tractable, we benchmarked multiple selection algorithms and introduced greedier forward selection, a linear-time algorithm that identifies near-optimal combinations with significantly reduced cost. This method enabled practical application of fusion without sacrificing performance

We then tested the real-world utility of this approach through two case studies: homologous sequence recovery and PPI prediction. In both settings, embedding fusion paired with greedier forward selection led to new SoTA results, outperforming previous baselines. These findings show that improved representations can substantially enhance model performance and practical utility. More broadly, they demonstrate how our approach transforms embedding fusion from a promising but computationally intractable idea into a usable tool for protein machine learning.

We also acknowledge the limitations of our study. Recent works have highlighted the species bias present in the training data for pLMs as well as the potential data leakage between the pre-training data and the downstream task [37,38]. As a result, performance with any pLM embeddings on benchmark datasets may be over optimistic in regard to its ability to generalize to unseen or underrepresented proteins. While datasets used in this study have been partitioned to consider the sequence similarity between the training and test splits, the pre-training data and test data are still relatively small and limited. As foundation models continue to scale and absorb increasingly diverse training data, future work should focus on evaluating representation robustness and transferability under more challenging generalization settings.

## Methods

### Protein language models

Table 1. provides an overview of selected tpLMs. We describe the key features of each tpLM, below. additional details, including the composition of pre-training data, are described in **Supplementary Section S1**.

OntoProtein utilized Gene Ontology and Gene Annotation terms during pre-training and showed improvements on the TAPE benchmarks [39], protein-protein interaction prediction, and protein function prediction [40]. ProtST performed contrastive learning between PubMedBERT embeddings of Swiss-Prot text annotations and pLM embeddings to achieve further improvements in downstream performance [41]. Similarly, ProteinDT introduced the Contrastive Language and Protein pre-training (ProteinCLAP) approach to align the representations between SciBert [42] embeddings of Swiss-Prot annotations and ProtBert [23] embeddings. ProteinCLIP used contrastive learning to align protein sequence representations from pre-trained pLMs to UniProt function descriptions embedded by OpenAI’s text embedding model [19]. The tri-modal pLM ProTrek was trained with a contrastive learning objective between sequence, structure, and function, aligning GPT-4-summarized UniProt descriptions embedded by PubMedBERT and protein sequence embedded by ESM2 [20]. Lastly, ESM3 takes a different tri-modal approach, representing sequence, structure, and function annotations as discrete tokens and predicting the masked tokens across the three different modalities as the pre-training task [43].

### Datasets

We adopt the same six benchmarks used by Schmirler et al.’s study on fine-tuning pLMs [30]. We exclude the disorder [44] and secondary structure prediction [45,46] tasks because protein structure tasks have already been shown to be well aligned with current pLM pre-training [18]. The datasets for homologous sequence recovery PPI prediction have previously been used by Wu et al. [19]. Detailed descriptions of datasets as well as dataset statistics are described in **Supplementary Section S2 and Supplementary Table 4**.

### Training and Implementation

All models used in this study are re-implementations of existing architectures [19,30,35], which are described below. Models are implemented in PyTorch v2.2.2 and trained on one NVIDIA RTX A6000 GPU with 48Gb memory. All deep-learning models are run five times using five different seeds (2,4,8,16, and 32), checkpointed based on the best validation performance, and the average performance on the test set across seeds is reported. Technical details regarding embedding generation, implementation of embedding fusion, and the metrics used in this study are described in **Supplementary Section S6 and S7**.

### Evaluation Metrics

For benchmarking tpLMs and embedding fusion, we measure two metrics: **Spearman’s** ρ (Spearman’s rank correlation coefficient) measures the monotonic relationship between the predicted and true values by comparing their rank orderings.

**Top-1 accuracy** is used for classification tasks where a single label is predicted for each input. It reflects the proportion of correct predictions among all samples and is used for the Location benchmark.

For the homologous sequence recovery task, we also use top-1 accuracy, where a retrieval is considered correct if the nearest-neighbor sequence (based on cosine similarity in embedding space) belongs to the same homologous superfamily as the query.

For the protein-protein interaction (PPI) prediction task, we report several standard metrics for binary classification:

#### Accuracy

The fraction of correct predictions across all positive and negative samples.

#### AUPRC (Area Under the Precision-Recall Curve)

Measures the trade-off between precision and recall across thresholds. It is especially informative under class imbalance, as it focuses on the performance for the positive class.

#### F1 Score

The harmonic mean of precision and recall, providing a balanced metric when false positives and false negatives carry similar costs.

#### Matthews Correlation Coefficient (MCC)

A balanced metric that accounts for all four values of the confusion matrix. It is considered a robust overall measure of binary classifier performance, particularly in imbalanced settings.

### Regressors and Classifier for the six benchmark datasets

The use of a shallow multilayer perceptron (MLP), referred to as the prediction head, is standard practice in recent pLM benchmarking studies, such as PEER [47], which use prediction heads to isolate the representational power of the pLM. This design allows for fair comparison across models and facilitates cross-referencing of results across studies.

We adopt the prediction head used by Schmirler et al. [30]. All models begin with batch normalization of the input data, followed by a hidden layer, ReLU activation, dropout, and an output layer. The output dimension is 10 for the subcellular location classifier and 1 for all other regressors. The loss function is Cross Entropy Loss for the subcellular location classifier and Mean Squared Error Loss for all other regressors. The loss is optimized via Adam without a learning rate scheduler. The hidden dimension size is set to 32 and the dropout rate is set to 20%. While all other hyperparameters are unchanged from Schmirler et al., we scaled the batch size from 8 to 4096 to reduce training time. We scaled the original learning rate from 0.0001 to 0.0001×√512 based on the guidance by Malladi et al. [48] to match the change in batch size. All hyperparameters are displayed in **Supplementary Table 5**.

### Embedding Selection Algorithms

#### Forward Selection

Forward selection begins with an empty set and iteratively adds the embedding that most improves performance when combined with the current set. At each step, all remaining embeddings are evaluated in combination with the current subset, and the one that yields the best validation performance is added. This process continues until no further improvement is observed or all embeddings have been evaluated. This algorithm is O(N^2^) in the worst case, due to repeated evaluation of all remaining candidates at each step.

Algorithm pseudocode:

1. Initialize selected set S = □
2. While improvement is observed:
  - For each e □ S, evaluate performance of S □ {e}
  - Add e* that gives the highest performance
3. Return best-performing subset S

#### Backward Selection

Backward selection starts with the full set of embeddings and iteratively removes the embedding whose removal most improves (or least degrades) performance. This algorithm is also O(N^2^), as each step evaluates all remaining removals.

Algorithm pseudocode:

1. Initialize selected set S = {all embeddings}
2. While performance does not degrade:
  - For each e □ S, evaluate performance of S \ {e}
  - Remove e* that yields the highest performance
3. Return best-performing subset S

#### Greedier Forward Selection

We begin by evaluating the performance of each individual embedding on the validation set and ranking them from best to worst. Let the ranked list be [e_1_,e_2_,…,e_N_], where e_1_ yields the highest standalone performance. Starting with the top-performing embedding {e_1_}, we iteratively evaluate the effect of adding the next-best ranked embedding to the current set. That is, we test {e_1_,e_2_}, then {e_1_,e_2_,e_3_}, and so on. At each step, an embedding is added only if it improves performance on the validation set; otherwise, it is skipped. This process continues until all embeddings have been considered. This method requires N evaluations for single-embedding ranking and up to N−1 additional evaluations for combinations, making it significantly more efficient than standard forward or backward selection. See **Supplementary Figure 1** for pseudocode.

### Homologous sequence recovery

We use the state-of-the-art homologous sequence recovery framework from Wu et al. [19]. Given a list of query proteins and a protein database, we calculate the cosine similarity between the query protein embedding and embedding of every protein in the database. We retrieve the protein with the highest similarity to the query, excluding itself, and check if the query protein and the retrieved protein belong to the same CATH superfamily. If they belong to the same homologous superfamily, we consider that a successful retrieval. The reported accuracy is the number of successful retrievals divided by the number of queries made. There are no hyperparameters for this retrieval pipeline. Following Wu et al., we L2-normalize all embeddings before the retrieval pipeline.

### Protein-protein interaction classifier

We use the state-of-the-art protein-protein interaction classifier used by NaderiAlizadeh and Singh [35]. The protein-protein interaction classifier uses one projection layer followed by a ReLU activation. The binary prediction (0 or 1) is determined by rounding the cosine similarity between the two protein projections at the 0.5 threshold. The model uses Binary Cross Entropy as the loss function, which is optimized using the AdamW optimizer paired with a Cosine Annealing Warm Restarts learning rate scheduler that restarts every 10 epochs. The hidden dimension size is set to 1024. The weights from the epoch with the highest validation AUPRC is used on the test set. These hyperparameters are the set by NaderiAlizadeh and Singh [35]. While we scaled the batch size from 32 to 4096, we kept the learning rate constant as no change in performance was observed. Exact hyperparameters are shown in **Supplementary Table 6**.

## Key Points

- No single protein language model representation consistently performs best across all biological tasks, highlighting a need for methods that can combine complementary strengths of different embeddings.
- Embedding fusion improves the quality of protein representations but the best combination is task-specific and hard to identify without the costly exhaustive search.
- We propose greedier forward selection, a fast and scalable algorithm that enables near-optimal embedding fusion, leading to new state-of-the-art results in homologous sequence recovery and protein-protein interaction prediction.

## Data and Code Availability

The data and code necessary to reproduce the results from this study are available at https://github.com/Wang-lab-UCSD/Benchmarking-tpLMs.

## Author Contributions

**Young Su Ko:** methodology, software, investigation, and writing (original draft). **Jonathan Parkinson:** software, validation, writing (review and editing). **Wei Wang:** supervision and writing (review and editing).

## Ethics declarations

The authors declare no competing interests.

## Author Information

**Young Su Ko** is a PhD candidate at the University of California, San Diego.

**Jonathan Parkinson** is a Project Scientist at the University of California, San Diego.

**Wei Wang** is a Professor at the University of California, San Diego.

