## Supplementary Information for "Scalable embedding fusion with protein language models: insights from benchmarking text-integrated representations"

**Section S1. Detailed descriptions of selected protein language models.**

#### **ESM2**

ESM2 is a standard pLM we use to compare against the tpLMs. ESM2 is trained at six different scales, ranging from 8M parameters to 15B parameters. We selected the 3B parameter model to serve as the baseline pLM for comparison.

**Ankh**

Ankh is another pLM trained using a MLM objective. We use the Ankh-large variant, which contains 1.9B parameters, as a second strong non-text baseline in our comparisons.

**ProtT5**

ProtT5 is a protein-adapted version of the T5 language model. It is trained using Bart-like MLM denoising objective on a diverse set of protein sequences from the BFD and UniRef50 datasets. In this study, we use the 3B parameter version of ProtT5, called ProtT5-XL-UniRef50.

#### **ProteinCLIP**

The text data for training ProteinCLIP originates from the UniProt **“Function”** section.

There are six variations of ProteinCLIP, where the base model is one of ProtT5, ESM2 8M, 35M, 150M, 640M, and 3B. To select the ProteinCLIP model for the six benchmark tasks, we calculate the average accuracy of the six models on ProteinCLIP’s reported tasks of protein-protein interaction prediction and homologous protein recovery. As ProteinCLIP trained on top of ProtT5 has the highest average accuracy, we use the ProtT5 variation for the six initial benchmark tasks. For our application of embedding fusion on the protein-protein interaction and homologous sequence recovery tasks we use the ProteinCLIP models with the best reported performance, ProtT5 and ESM2 3B respectively.

#### **ProteinDT**

The SwissProtCLAP text-protein pair dataset constructed by the authors of ProteinDT uses the SwissProt annotated subset of UniProt. From SwissProt, the **“Function”**, **“PTM/Processing”**, and **“Interaction”** sections are used.

ProteinDT offers two variations ProteinDT-InfoNCE and ProteinDT-EBM-NCE, differing in the loss function used during pre-training. We select ProteinDT-InfoNCE as it has a higher average performance across the benchmarks tested by the authors.

#### **ProtST**

ProtST uses text from the SwissProt annotated subset of UniProt. From SwissProt, the **“Function”**, **“Subcellular Location”**, and **“Sequence similarities”** sections (under “Family & Domains”).

ProtST provides three options: ProtST-ProtBert, ProtST-ESM-1b, and ProtST-ESM2, differing in the base model used. Based on the performance on function annotation and the recommendation provided by the authors, we select ProtST-ESM-2 for this study.

#### **ProTrek**

ProTrek uses text from the UniProt database. The authors state that “**almost all subsections**” of the UniProt database are used.

ProTrek offers two pre-trained models with different parameter sizes (35M and 650M). We select the 650M model as it was highlighted by the authors in their comparison with other state-of-the-art pLMs.

#### **OntoProtein**

OntoProtein pairs **Gene Ontology** and **Gene Annotations** with protein sequence.

There is only one pre-trained OntoProtein model available.

#### **ESM3**

ESM3 uses residue-level text descriptions from InterPro annotations (annotation term names, associated GO terms per annotation, and ancestor GO terms).

There is only one pre-trained ESM3 model available.

**Section S2. Details of datasets used in study.**

### **GB1**

The GB1 dataset (1) is a mutational dataset measuring the fitness (stability and binding affinity) of different mutants of the binding domain of Protein G. We use the 3-vs-rest split created by the FLIP benchmark (2) authors. The training and validation data consist of wild type, single, double and triple mutants, while the test data consist of all other mutants.

#### **GFP**

The GFP dataset (3) is a deep mutational scanning dataset that reports the fluorescence intensity of different green fluorescent protein mutants. The training, validation, and test splits are created by the TAPE benchmark (4) authors. The training and validation data are in a Hamming distance 3 neighborhood of the original sequence, while the test data is from the Hamming distance 4-15 neighborhood.

#### **AAV**

The AAV dataset (5) is a mutational screening dataset that measures the fitness of different adeno-associated virus 2 capsid protein mutants. We use the 2-vs-rest split created by the FLIP benchmark (2) authors. The training and validation data consists of wild type, single, and double mutants while the test data consists of all other mutants.

#### **Location**

The Location dataset (6,7) is a 10-class classification dataset that involves the prediction of a protein’s subcellular location. The training and validation originates from DeepLoc (6) while the test set is Stärk et al.'s (7) “setHard” dataset, in which no sequences have more than 20% pairwise sequence identity from the DeepLoc training and validation data. The “setHard” test set provides a better evaluation of a pLM’s learned representation on unseen data than the DeepLoc test set.

#### **Meltome**

The Meltome dataset (8) collects the melting temperatures of proteins from 13 species. We use the mixed split created by the FLIP benchmark (2) authors. After all sequences are clustered, cluster representatives are selected using MMseqs2 at a 20% sequence identity threshold. Training data consists of sequences in 80% of clusters while the test data only consists of cluster representatives in the remaining 20% of clusters.

#### **Stability**

The Stability dataset (9) collects protease susceptibility measurements. We use the split created by the TAPE benchmark (4) authors. The training and validation data consists of sequences from four rounds of experimental tests while the test data consists of sequences in 1-Hamming distance neighborhoods around the promising proteins observed in the four experimental rounds.

#### **Protein-Protein Interaction Prediction**

The “gold-standard” human protein-protein interaction dataset created by Bernett et al. (10) is a binary classification dataset (interacting pair vs non-interacting pair). There is no overlap between training, validation, and test datasets. In addition, the sequence identity of sequences across sets are capped at 40%.

#### **Homologous Sequence Recovery**

The CATH dataset (11) consists of protein domains. Following Wu et al. (12) we use version 4.3.0 and use CATH S20, in which two sequences have a maximum sequence identity of 20% and 60% overlap. All proteins in non-singleton superfamilies are designated as query proteins. There is no training, validation, and test split for this dataset as it is not used for a machine learning task.

**Section S3. Normalization for input dimension size.**

Recognizing that embedding fusion increases the input dimension, and thus the parameter count in the projection layer, we implemented a control procedure. Specifically, for each benchmark, after identifying the best combined embedding, we concatenated the best-performing single embedding multiple times to itself to match the input dimension of the combined embedding. If the input dimension could not be matched exactly, we rounded up, giving the repeated single embedding a slight advantage. If this increased input dimension reduced performance, we reverted to the original single embedding for comparison. The best embedding combinations and their matched single embedding baselines for each benchmark are detailed in **Supplementary Table 3**.

**Section S4: Effects of increasing model size on memory and runtime.**

An important area of consideration is the increased memory and runtime incurred by the increase in input dimensionality from embedding fusion. In our benchmark experiments, we observed an average increase in runtime by 1.65x. Although the absolute increase in runtime was always less than a minute, a concern is that in bigger models, the increase in runtime will scale linearly. For example, a model that takes 100 minutes to train may become 165 minutes. However, we reasoned that the majority of the runtime is incurred early on in the fusion, such as the projection layer, and bigger models do not necessarily incur a proportional slow-down.

To demonstrate, we use the GB1 benchmark, as the best combination for this benchmark had the highest-dimensional fused embedding (5258 dimensions). We train two models: the original architecture (178,497 parameters), and a wider model with layers [5258 → 3200 → 32 → 1], totaling 1.7 million parameters–a 9.5× increase in model size.

Despite this 9.5× increase in parameters, the maximum GPU memory usage and runtime increased by less than 2×, illustrating that embedding fusion remains efficient even in larger networks. This demonstrates that absolute costs, not just relative ratios, must be considered when assessing scalability, and that the cost of fusion does not explode with model size. Therefore, we believe these overheads do not meaningfully limit the practical adoption of embedding fusion, even in large-scale protein modeling pipelines.

| Model | Parameter Count | Max GPU Memory (Mb) | Runtime (s) |
| --- | --- | --- | --- |
| Original | 178497 | 181.3804 | 20.66462 |
| Scaled-up Model | 1700481 | 217.0405 | 34.13523 |
| Ratio | 9.52 | 1.19 | 1.65 |

*Max GPU Memory and runtime are averaged over 5 random seeds.

**Section S5. Reproducing NaderiAlizadeh PPI Classifier**

We reproduce the state-of-the-art PPI classifier, as the pre-trained model checkpoints are not available. Using the same hyperparameters, we are able to reproduce nearly identical results, falling within the standard deviation of each other. We report the mean and standard deviation across 5 seeds.

| Model | Test PRC | Test Accuracy | Test Sensitivity | Test Specificity | Test Precision | Test F1 | Test MCC |
| --- | --- | --- | --- | --- | --- | --- | --- |
| Reported | **0.717±0.001** | 0.657±0.002 | **0.696±0.022** | 0.618±0.024 | 0.646±0.007 | **0.670±0.006** | 0.315±0.003 |
| Ours | 0.716±0.002 | **0.658±0.004** | 0.679±0.034 | **0.637±0.042** | **0.652±0.014** | 0.665±0.008 | **0.317±0.007** |

### **Section S6. Embedding generation process.**

To generate the embeddings, we follow the instructions provided by the respective authors in their GitHub repositories. For six out of our nine tested pLMs, an N-amino acid long protein sequence input returns a [start token + N + end token, hidden dimension] shaped representation. We first remove the start and end tokens then mean-pool across the length axis to return a fixed-length vector for each protein. For ProtST, ProteinCLIP, and ProTrek, this step is not necessary as the model by default returns a fixed length vector.

Due to memory constraints, we set the maximum protein sequence length to be 5800AA. For sequences longer than 5800AA, we select the first 2400 residues and the last 2400 residues, concatenate the residues to create a pseudo-protein sequence following Li et al. (16) who reasoned taking the N and C terminuses preserves biologically relevant signals.

**Section S7. Technical details of implementing embedding fusion.**

To create a combined embedding, we concatenate the fixed length representations as a preprocessing step. Each protein language model is designated by an alphanumeric character (0: Ankh, 1: ESM2, 2: ProtT5, B: ESM3, C: OntoProtein, D: ProteinCLIP, E: ProtST, F: ProTrek, G: ProteinDT). For six different tpLM embeddings, we explore 2^6^-1=63 possible combinations–subtracting one to exclude the empty set. We do not consider the order of the embeddings, thus combinations are always sorted alphabetically by their one-letter code. For instance, the combination of embeddings G, D, and E would be ordered as {DEG} in the concatenated representation.

**Supplementary Table 1.** Mean and 95% confidence intervals for six benchmark datasets.

| Model | AAV | GB1 | GFP | Location | Meltome | Stability |
| --- | --- | --- | --- | --- | --- | --- |
| ESM2 3B | **0.770±0.201** | 0.833±0.004 | 0.613±0.002 | 0.627±0.011 | 0.663±0.004 | 0.733±0.015 |
| Ankh | 0.681±0.005 | 0.838±0.004 | 0.647±0.002 | 0.598±0.016 | 0.590±0.013 | 0.705±0.034 |
| ProtT5 | 0.690±0.013 | **0.844±0.007** | 0.595±0.003 | 0.609±0.014 | 0.674±0.003 | 0.766±0.021 |
| OntoProtein | 0.671±0.009 | 0.737±0.011 | **0.666±0.002** | 0.542±0.007 | 0.648±0.008 | 0.677±0.023 |
| ProteinDT | 0.543±0.032 | 0.800±0.007 | 0.573±0.003 | 0.563±0.011 | 0.447±0.006 | 0.610±0.042 |
| ProtST | 0.621±0.010 | 0.815±0.007 | 0.553±0.002 | 0.640±0.014 | 0.550±0.008 | 0.699±0.036 |
| ProteinCLIP | 0.546±0.019 | 0.790±0.002 | 0.469±0.005 | 0.613±0.005 | 0.420±0.006 | 0.601±0.033 |
| ProTrek | 0.344±0.028 | 0.837±0.004 | 0.645±0.001 | **0.737±0.010** | **0.689±0.005** | 0.597±0.047 |
| ESM3 | 0.641±0.010 | 0.818±0.011 | 0.640±0.002 | 0.561±0.007 | 0.599±0.007 | **0.779±0.035** |

For **all subsequent tables**, we assign an alphabetic identifier for each model for readability.

| ESM3 | OntoProtein | ProteinCLIP | ProtST | ProTrek | ProteinDT |
| --- | --- | --- | --- | --- | --- |
| B | C | D | E | F | G |

**Supplementary Table 2.** Test set performances of single tpLMs and combination of tpLMs across six benchmark datasets.

| Dataset | Best Single |  | Best Single Repeated |  | Best Combination |  |
| --- | --- | --- | --- | --- | --- | --- |
| AAV | C (1024) | 0.671±0.009 | CCCC (4096) | 0.647±0.031 | BCDE (3210) | **0.732±0.006** |
| GB1 | F (1024) | 0.837±0.004 | FFFFFF (6144) | 0.831±0.005 | BCDEFG (5258) | **0.849±0.007** |
| GFP | C (1024) | 0.666±0.002 | CC (2048) | 0.664±0.005 | CF (2048) | **0.671±0.001** |
| Location | F (1024) | **0.737±0.010** | FF (2048) | 0.722±0.008 | DF (1152) | 0.733±0.007 |
| Meltome | F (1024) | 0.689±0.005 | FFFFFF (6144) | 0.700±0.004 | BCEFG (5130) | **0.728±0.006** |
| Stability | B (1546) | 0.779±0.035 | BB (3092) | 0.783±0.042 | BDF (2968) | **0.804±0.033** |

In parentheses, we denote the size of each input embedding. To account for the fact that combined embeddings will typically have a larger input dimension size, resulting in a projection layer with more parameters, it is possible that performance gains observed from embedding fusion are actually a consequence of the increased parameter count. To check, we trained models in which we combined the single embedding multiple times to match the number of parameters of the best combination. In cases it cannot be exactly matched, the repeated embeddings were given an advantage with more parameters. We see that in many cases, repeating reduced performance. Even under the parameter-normalized conditions, the improvements from embedding fusion remain significant.

**Supplementary Figure 1.** Pseudocode for optimal embedding combination search.

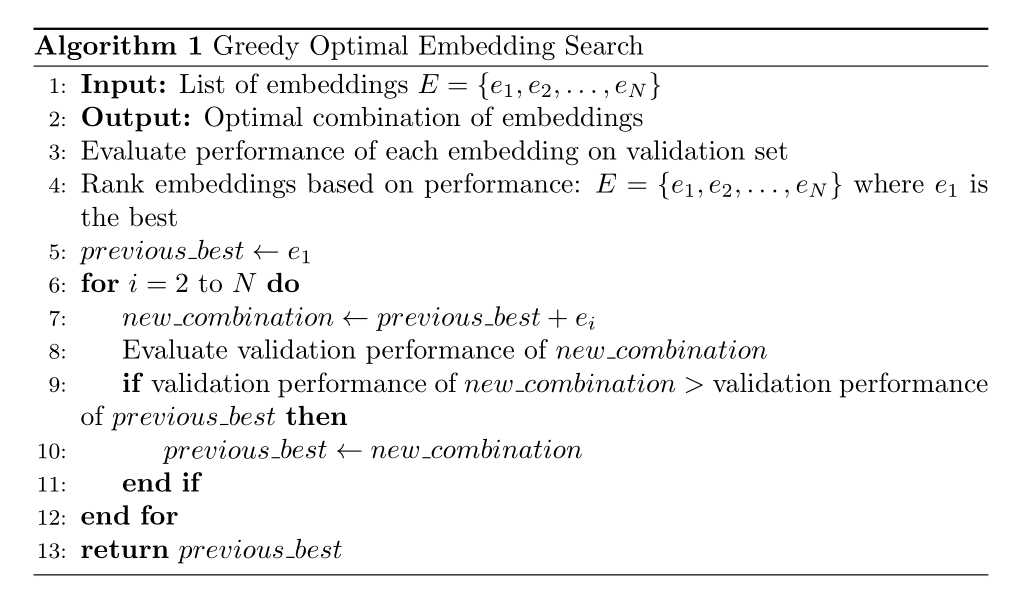

**Supplementary Table 3.** Application of greedier forward selection for protein-protein interaction prediction.

| Embedding | AUPRC* | Iteration | Current Best | AUPRC | Potential Best | AUPRC | Combine? |
| --- | --- | --- | --- | --- | --- | --- | --- |
| ProtST | 0.696 | 1 | E | 0.696 | DE | **0.708** | Yes |
| ProteinCLIP | 0.692 | 2 | DE | 0.708 | DEF | **0.714** | Yes |
| ProTrek | 0.691 | 3 | DEF | **0.714** | DEFG | 0.709 | No |
| ProteinDT | 0.672 | 4 | DEF | **0.714** | BDEF | 0.664 | No |
| ESM3 | 0.663 | 5 | DEF | **0.714** | CDEF | 0.712 | No |
| OntoProtein | 0.660 | **Return** | **DEF** |  |  |  |  |

*Single embedding performance is shown in this column, all of which are outperformed by the identified combinations shown in Figure 4G of the main text.

**Supplementary Table 4.** Details of datasets used.

| Dataset | Train | Validation | Test | Average Length |
| --- | --- | --- | --- | --- |
| AAV | 28,626 | 3,181 | 50,776 | 736.3 |
| GB1 | 2,691 | 299 | 5,743 | 265 |
| GFP | 21,446 | 5,362 | 27,217 | 237.0 |
| Location | 9,503 | 1,678 | 490 | 519.9 |
| Meltome | 22,335 | 2,482 | 3,134 | 544.5 |
| Stability | 53,614 | 2,512 | 12,851 | 45.0 |
| Protein-protein* | 163,192 | 59,260 | 52,048 | 630.1 |
| Dataset | Query | Database |  | Average Length |
| CATH S20 | 10,926 | 14,941 |  | 155.2 |

*The protein-protein dataset is balanced, with an equal number of positive and negative examples.

**Supplementary Table 5.** Hyperparameters for six benchmark datasets.

| Dataset | Hidden Dimensions | Epochs | Learning Rate | Batch Size | Adam Epsilon |
| --- | --- | --- | --- | --- | --- |
| AAV | 32 | 120 | 0.00226274169 | 4096 | 1e-7 |
| GB1 | 32 | 240 | 0.00226274169 | 4096 | 1e-7 |
| GFP | 32 | 240 | 0.00226274169 | 4096 | 1e-7 |
| Location | 32 | 120 | 0.00226274169 | 4096 | 1e-7 |
| Meltome | 32 | 120 | 0.00226274169 | 4096 | 1e-7 |
| Stability | 32 | 120 | 0.00226274169 | 4096 | 1e-7 |

**Supplementary Table 6.** Hyperparameters for protein-protein interaction prediction.

| Hyperparameter | Value |
| --- | --- |
| Hidden Dimensions | 1024 |
| Epochs | 50 |
| Learning Rate | 0.0001 |
| Batch Size | 4096 |
| Epochs before CosineAnnealing Restarts | 10 |

**Supplementary Table 7.** Test set performances of heuristic-identified combinations and true optimum combinations on six benchmark datasets.

| Dataset | Heuristic-identified | Performance | True Optimum | Performance |
| --- | --- | --- | --- | --- |
| AAV | BCDE | 0.732±0.006 | BCDE | 0.732±0.006 |
| GB1 | BDEFG | 0.837±0.007 | BCDEFG | 0.849±0.007 |
| GFP | CF | 0.671±0.001 | CF | 0.671±0.001 |
| Location | EF | 0.724±0.006 | F | 0.737±0.010 |
| Meltome | BCDEFG | 0.725±0.003 | BCEFG | 0.728±0.006 |
| Stability | BC | 0.755±0.030 | BDF | 0.804±0.033 |

**Supplementary File 1.** Excel file containing the training logs of all models used in this study.
